# Locomotion-induced ocular motor behavior in larval *Xenopus* is developmentally tuned by visuo-vestibular reflexes

**DOI:** 10.1101/2021.01.04.424105

**Authors:** Julien Bacqué-Cazenave, Gilles Courtand, Mathieu Beraneck, Hans Straka, Denis Combes, François M. Lambert

## Abstract

Locomotion requires neural computations to maintain stable perception of the world despite disturbing consequences of the motor behavior on sensory stability. The developmental establishment of locomotor proficiency is therefore accompanied by a concurrent maturation of gaze-stabilizing motor behaviors. Using developing larval *Xenopus*, we demonstrate mutual plasticity of predictive spinal locomotor efference copies and multi-sensory motion signals with the aim to constantly ensure dynamically adequate eye movements during swimming. Following simultaneous ontogenetic onsets of locomotion, spino-ocular, optokinetic and otolith-ocular motor behaviors, locomotor efference copy-driven eye movements improve through gradually augmenting influences of semicircular canal signals. Accordingly, neuronal computations change from a predominating cancelation of angular vestibulo-ocular reflexes by locomotor efference copies in young larvae to a summation of these signals in older larvae. The developmental switch occurs in synchrony with a reduced efficacy of the tail-undulatory locomotor pattern generator causing gradually decaying influences on the ocular motor output.

## INTRODUCTION

Locomotion requires concurrent neural computations that optimally maintain sensory perception despite the disturbing consequences of the motor behavior on the encoding and transformation process. Accordingly, all vertebrates optimize their visual acuity by integrating feedback signals from passively-induced head/body motion with intrinsic feedforward signals during self-generated movements^1, 2, 3^. Motion-induced retinal image displacements are minimized by counteractive eye and/or head-adjustments that derive from the synergistic activity of visual, vestibular and proprioceptive reflexes^3, 4, 5^, supplemented during self-motion by efference copies of the propulsive motor commands^6, 7^. Independent of the locomotor style, such feedforward replica from spinal pattern-generating circuits evoke spatio-temporally adequate eye movements as originally demonstrated in pre- and post-metamorphic amphibians^8, 9, 10^. Suggestive evidence for a ubiquitous role of such signals in all vertebrates was subsequently provided by studies in mice^11^ and human subjects^12, 13^. Gaze-stabilizing eye movements during locomotion, therefore result from an integration of locomotor efference copies and motion-related sensory signals into ocular motor commands. The computational algorithms for integrating these signals are likely established prior to and during the developmental acquisition of locomotion and further modified as propulsive proficiency increases.

Progressive developmental improvements of locomotor performance are tightly linked to the maturation of gaze-and posture-stabilizing reflexes^14, 15, 16^. The assembly and functional initiation of inner ear structures and associated sensory-motor circuitries are decisive factors for the onset and efficiency of gaze-stabilization and locomotion^17^. This is particularly critical in small aquatic vertebrates, where coordinated locomotor activity is generally acquired very early after hatching^14, 17^. In accordance with this requirement, utriculo-ocular reflexes become functional immediately after hatching in zebrafish^18^ and anuran larvae^19^ followed, though conspicuously later, by semicircular canal-derived vestibulo-ocular reflexes (VOR)^15, 20^. The different developmental acquisition of individual VOR components, as demonstrated in *Xenopus* larvae, but likely also of visuo-ocular reflexes, might have significant interactive influences on the maturation of locomotor proficiency with respective reciprocal consequences for efference copy-driven gaze-stabilization^8^. Since swim-related head motion in e.g., *Xenopus* tadpoles represents a reliable sensory reference, vestibular signals might be well-suited to guide the ontogenetic progression of locomotion-induced ocular motor behaviors.

Here, we demonstrate the mutual developmental plasticity of spinal locomotor efference copies and multi-sensory motion signals with the aim to produce dynamically adequate gaze-stabilizing eye movements during swimming in larval *Xenopus*. Following simultaneous onsets of locomotor activity, spino-ocular, optokinetic and otolith-ocular motor behaviors, efference copy-evoked predictive eye movements gradually improve through the influence of semicircular canal signals during larval development until metamorphic climax. During this period, the sensory-motor integration changes from a cancelation of the angular VOR by locomotor efference copies in young larvae to a summation of the respective signals in older larvae. The developmental switch in computational outcome occurs in synchrony with a reduced efficacy of the tail-undulatory locomotor pattern generator causing a gradually decaying influence on the ocular motor output.

## MATERIALS AND METHODS

### Animals

Experiments were conducted on larvae of the South African clawed toad *Xenopus laevis* of either sex (*N* = 149), obtained from the Xenopus Biology Resources Centre (University of Rennes 1, France; http://xenopus.univ-rennes1.fr/). Animals were maintained at 20–22 °C in filtered aquarium water with a 12:12h light/dark cycle. Experiments were performed on tadpoles between developmental stage 40 and 58, classified by stage-specific morphological features^21^. All procedures were carried out in accordance with and approved by the local ethics committee (#2016011518042273 APAFIS #3612).

### Video recordings and analysis of eye and tail movements in semi-intact preparations

The performance of ocular motor behaviors during passively imposed head movements, visual image motion and undulatory swimming was recorded and quantified in semi-intact preparations^9^. To generate such preparations, tadpoles were anesthetized in an aqueous solution of 0.05% 3-aminobenzoic acid ethyl ester methanesulfonate (MS-222) and subsequently transferred to a dish containing an oxygenated (95% O_2_, 5% CO_2_) Ringer solution (NaCl, 120 mM, KCl, 2.5 mM, CaCl_2_ 5 mM; MgCl_2_ 1 mM, NaHCO_3_ 15 mM; pH 7.4) at ∼18°C. Isolation of the preparation was achieved by removal of the lower jaws and evisceration or decapitation at various, experiment-dependent levels of the spinal cord. The central nervous system, inner ears, eyes and motor effectors including efferent and afferent neuronal innervation, respectively, remained functionally preserved ^22^. Following removal of the skin, the cartilaginous skull was opened from dorsal to disconnect the forebrain and to expose the hindbrain thereby providing access for the Ringer solution during the submerged placement in the recording chamber. The preparation was firmly secured with the dorsal side up to a Sylgard floor and continuously superfused with oxygenated Ringer solution at a rate of 1.5 - 2.0 ml/min.

### Recordings of angular and gravitoinertial vestibulo-ocular reflexes

Horizontal and vertical eye movements were elicited during application of head motion stimuli. For this purpose, the recording chamber was mounted onto a computer-controlled, motorized turn-table (Technoshop COH@BIT, IUT de Bordeaux, Université de Bordeaux). The semi-intact preparation in the chamber was positioned such that the vertical rotation axis was centered in the midline between the bilateral otic capsules and the horizontal axis was located at the same dorso-ventral level as the otic capsules to ensure optimal activation of all semicircular canals and the utricle. Horizontal head motion stimulation consisted of sinusoidal rotations around the vertical axis at a frequency of 1 Hz and maximal stimulus position eccentricity that ranged from ± 5° - 25°. Head roll motion stimulation consisted of sinusoidal rotations around a rostro-caudally directed horizontal axis at a frequency of 0.25 Hz and maximal stimulus position eccentricity that ranged from ± 2.5° - 20°. To exclude contributions of visuo-motor reflexes and locomotor efference copy-driven eye movements to the responses evoked by imposed head motion in particular subsets of the experiments, both optic nerves were transected at the level of the optic chiasm and the tail was severed at the level of the first spinal segments to eliminate ascending spino-ocular motor commands^8^.

### Recording of optokinetic reflexes

Horizontal visual motion-induced eye movements were evoked by a rotating large-field visual scene. Stimulation was provided by placing the recording chamber with the semi-intact preparation in the center of a custom-made servo-controlled motorized drum with a diameter of 8 cm^23^. The wall of the rotating drum was illuminated from below, by visible light LEDs and displayed a pattern of equally spaced, vertical, black and white stripes with a spatial size of 15°/15°. The pattern was sinusoidally oscillated at a frequency of 0.25 Hz and maximal positional eccentricity of ± 2.5° - 20°.

### Recording of eye movements during undulatory tail motion

Horizontal eye movements were recorded during spontaneous episodes of left-right tail undulations^8^. Semi-intact preparations with intact tails were firmly secured by tight mechanical fixation of the head to the Sylgard floor of the recording chamber while the tail remained entirely mobile to execute swim movements. The strict mechanical immobilization of the head prevented any residual head movements and thus the activation of vestibular reflexes, while potential optokinetic reflexes were excluded by transection of both optic nerves (see above). Movements of the eyes and the tail during swimming and vestibular motion stimulation were video-recorded non-invasively at 500 frames per second (fps) with a high-speed digital camera (Basler AG, ac1920) equipped with a micro-inspection lens system (Optem MVZL macro video zoom lens, QIOPTIQ). For the acquisition of horizontal eye movements, the camera was placed above the preparation; vertical eye movements during head roll motion (gVOR) where acquired with the camera positioned sideward to one eye. Tracking of the eyes and tail was automated using a custom-made software, programmed in Python 3.5 environment^9^. The software performed a frame-by-frame calculation of the angles between the major axis of the oval-shaped eye and the longitudinal axis of the head, or the angle created by the positional deviation of the first five tail myotomes relative to the longitudinal axis of the head^9^.

### Recording of extraocular and spinal motor nerve spike discharge

Spike activity of ocular motor nerves and spinal ventral roots (Vr) were simultaneously recorded in semi-intact preparations as described previously^8, 24^. Respective preparations were produced as indicated above with the exception that the tail was not transversally severed but carefully dissected to expose the spinal cord along with the ventral roots until segment 22-25, while the adjoining distal part of the tail until the end was anatomically preserved. Selected Vrs along the dissected spinal cord were identified for the recordings. The *lateral rectus* (LR) eye muscle-innervating abducens nerve was disconnected from its target muscle close to the neuro-muscular innervation site, cleaned from connective tissue and prepared for the recording^8^.

Extracellular spike discharge of the LR motor nerve and the Vrs was recorded with glass suction electrodes, connected to a differential AC amplifier (A-M System Model 1700 AC, Carlsborg, US). Following production of the electrodes with a horizontal puller (P-87 Brown/Flaming, Sutter Instruments Company, Novato, CA, USA), tips were broken and individually adjusted to fit the LR nerve or Vr diameters. Electrophysiological signals were digitized at 10 kHz (CED 1401, Cambridge Electronic Design, UK), displayed and stored on a computer for offline analysis. In some experiments, recordings of the LR nerve of one eye were coupled to video recordings of the movements of the other eye. In this experimental setting, eye movements were captured with a Basler camera (specification see above), placed above the eye and were tracked using custom-made software as described recently^9^.

### Injection of hyaluronidase into the bilateral otic capsules

The formation of semicircular canals during development was prevented according to a previously established method^25, 26^. A single injection of a calibrated volume (10 nl) of the enzyme hyaluronidase (0.5 mg/mL, Sigma-Aldrich, France), dissolved in artificial endolymph solution was made into both otic capsules in larvae at developmental stage 44. Thereafter, animals were allowed to further develop until usage in experiments on semi-intact preparations. As demonstrated, such tadpoles anatomically lack all semicircular canals as well as corresponding ocular motor reflexes, while the utricle and associated sensory-motor computations remain functionally preserved^15, 25^.

### Signal processing and data analysis

Raw data of video or electrophysiological recordings were processed off-line and analyzed with Dataview v11.5.1 (by W.J. Heitler, University of St Andrews, Scotland). For video image processing, traces of movements obtained from the eyes and/or tail were first filtered with a 20 Hz low-pass filter. Maximal excursions of the eyes were identified for each individual motion cycle and used to calculate the amplitude of eye movements. Tail-based swim cycles were defined by the repetitive angular excursion of the tail, with deviations from the null angle (longitudinal head/body axis) as indicator for the onset of each cycle. Maximal angular excursions of tail and eyes (peak to peak amplitudes) were determined for each individual swim cycle and used to calculate instantaneous peak frequency and the temporal relationship between tail and eye movements. Head motion-evoked eye movement components during concurrent vestibular stimulation and fictive swimming were obtained by interpolating between each swim cycle-related average eye excursion. This procedure revealed the locomotion-triggered eye motion magnitude and dynamics, which would be otherwise masked by the vestibulo-ocular response component. Variations of eye motion amplitude/position during head motion were calculated for a minimum of four vestibular stimulus cycles. Traces with obvious stimulus-unrelated slow positional deviations were excluded from further analysis.

Spike discharge recordings of LR motor nerves and spinal Vrs were rectified and filtered (moving average, time constant: 25 ms) to obtain the integral of the spike activity. Temporal parameters of fictive swim episodes were obtained from the timing of the spinal Vr burst discharge. An event was marked for each peak detected in the Vr and LR recordings indicating the phase relation between spinal and ocular motor spike activity during each swim cycle. Frequency and temporal relationships were subsequently calculated using Dataview software.

### Spectral analysis

The principal frequency content of eye movements and LR motor nerve activity, corresponding to fictive swimming *versus* vestibular stimulation was extracted by spectral analyses. The task was performed with R software (v3.5.2, R Core Team, 2014; http://www.R-project.org/) and a custom-built R script from “wavelettComp” package (Roesch & Schmidbauer, 2014; Computational Wavelet analysis; R package v1.0; http://CRAN.R-project.org/package=WavelettComp)^27^. The spectral analysis consisted of two successive procedures, with a Fourier transformation (FFT) as first step, followed by a wavelet analysis. The FFT decomposes the recorded signal in multiple sinusoidal signals with different frequencies thereby identifying the principal oscillation frequencies. Thereafter, the wavelet analysis confirmed the principal oscillation frequencies and reconstructed sinusoidal signals corresponding to these frequencies. The range of oscillation frequencies usually ranged from 0.5 to 16 Hz. These results were presented as spectrograms representing the wavelet power spectrum of each continuous oscillation within the range of frequencies present in the recordings. Periodograms illustrate in addition the average power of each oscillation frequency. This required to down-sample the original spike discharge recording to 500 Hz (performed with Dataview software, interpolation method) in order to increase the computing velocity of the analysis algorithm.

### Computational model

A computational neural network model was created with the NEURON 7.3 program 28. Swim-related central pattern generator (swim CPG) circuitry with alternating rhythmicity consisted of bilateral pacemaker interneurons (cIN) coupled across the midline *via* a reciprocal inhibitory connection with and an individual excitatory interneuron (iEX) terminating on each cIN. The resultant activity is mediated onto output neurons (Sp output). Fictive swimming was initiated by a current pulse applied to a iEX. To mimic spino-ocular motor coupling, an efferent interneuron (EC) received a copy of the signal mediated onto the ipsilateral spinal output neuron to excite a contralateral abducens motoneuron (Abd Mot). Two distinct vestibular pathways were created by distinct neurons to simulate vestibulo-ocular (VO) and vestibulo-spinal (VS) pathways. These interneurons were individually recruited by a sinusoidal current with an adjustable intensity and frequency to mimic head rotation-related vestibular signals. A vestibulo-ocular reflex was simulated by exciting the VO interneuron, which in turn recruits a contralateral Abd Mot. The vestibulo-spinal pathway was modeled by activating a VS neuron, which in turn excites the ipsilateral spinal output neuron. Individual parameters were specified in material availability.

### Statistics

After signal processing in Dataview, all data sets were analyzed with Prism7 (GraphPad, USA) and results were expressed as mean ± standard error of the mean (SEM), unless stated otherwise. Differences between two data sets were tested for significance using the unpaired two-tailed Mann–Whitney *U*-test, and Kolmogorov-Smirnov test to compare distributions. Multiple values were compared with the non-parametric Kruskal-Wallis test followed by a Dunn’s multiple comparisons test. Correlations (*R*, Pearson correlation) and linear regression (*r*^*2*^) was used to evaluate correlations between eye and tail movement amplitudes during fictive swimming, or between the ratio of eye/tail movements and tail excursion amplitudes with the latter as indicator for the strength of swimming. Differences between data sets were assumed to be significant at *p* < 0.05.

The temporal relationships between eye and tail movements at different developmental stages were assessed by a circular phase analysis of pooled data (Oriana 4.02 software (Kovach Computing Services, Wales). The mean vector ‘*μ*’ and its length ‘*r*’ indicated the preferred phase and strength of coupling, respectively. For non-uniform distributions (tested with the Rayleigh’s uniformity test, *p*), mean phase values of each animal were plotted as the grand mean of individual means of phase relationships between eye and tail movements, expressed as (*µ, r, p*). The preferred direction of the grand mean vector was tested by Moore’s Modified Rayleigh test and the difference of phase relations between grand means at different developmental stages was evaluated by the Watson-Williams *F*-test.

## RESULTS

### Common onset of predictive, visual motion and otolith organ-evoked eye movements

Free swimming in *Xenopus laevis* larvae commences around stage 37/38 followed by a rapid acquisition of the typical kinematic profile that remains essentially unaltered between stage 38-40^28^ and stage 50^9^. The early developmental occurrence of undulatory swimming, however, is at variance with the lack of locomotor efference copy-driven gaze stabilizing spino-ocular motor behavior^8^ at stage 40 (Fig. 1a_1;_ Supplemental fig. 1b, *N* = 3). In fact, the earliest activation of reliable and robust compensatory eye movements triggered by ascending copies of spinal locomotor signals was observed at stage 42 and had a gain of 0.34 ± 0.08 (*N* = 3; Fig. 1a_2_; Supplemental fig. 1b). This onset coincided with the developmental acquisition of sensory stimulus-driven visuo- and gravitoinertial (otolith organ) ocular motor behaviors. In fact, stage 40 larvae were incapable of expressing a horizontal optokinetic (OKR; Fig. 1b_1;_ Supplemental fig. 1b, *N* = 5) or a gravitoinertial vestibulo-ocular reflex (gVOR; Fig. 1c_1;_ Supplemental fig. 1b, *N* = 5). First evidence for the presence of these sensory-motor-driven gaze stabilizing eye movements was encountered at stage 42 (Fig 1b_2_,c_2_). At that stage, the OKR and gVOR exhibited small, yet reliable eye movements with gains that exceeded 0.15 (0.16 ± 0.02 for OKR, *N* = 5 and 0.18 ± 0.05 for gVOR, *N* = 7; Supplemental fig. 1b). Thereafter, all gaze stabilizing eye movement components increased further in robustness to reach gains at stage 45 (*N* = 4) of 0.65 ± 0.02 for the spino-ocular motor coupling, 0.33 ± 0.08 for the OKR and 0.47 ± 0.21 for the gVOR (Fig. 1a_3_-c_3_, Supplemental fig. 1b). The performance continued to improve until maximal gain values were reached at stage 48 with a generally better performance of the gVOR compared to the OKR (Supplemental fig. 1c-e). The common developmental onset of all three gaze stabilizing eye movement components coincided with a major step in the maturation of the cyclic left-right spinal locomotor output, which changes from single-spikes at stage 37/38 to spike bursts at stage 42 (Supplemental fig. 1a)^29^.

**Figure 1:**
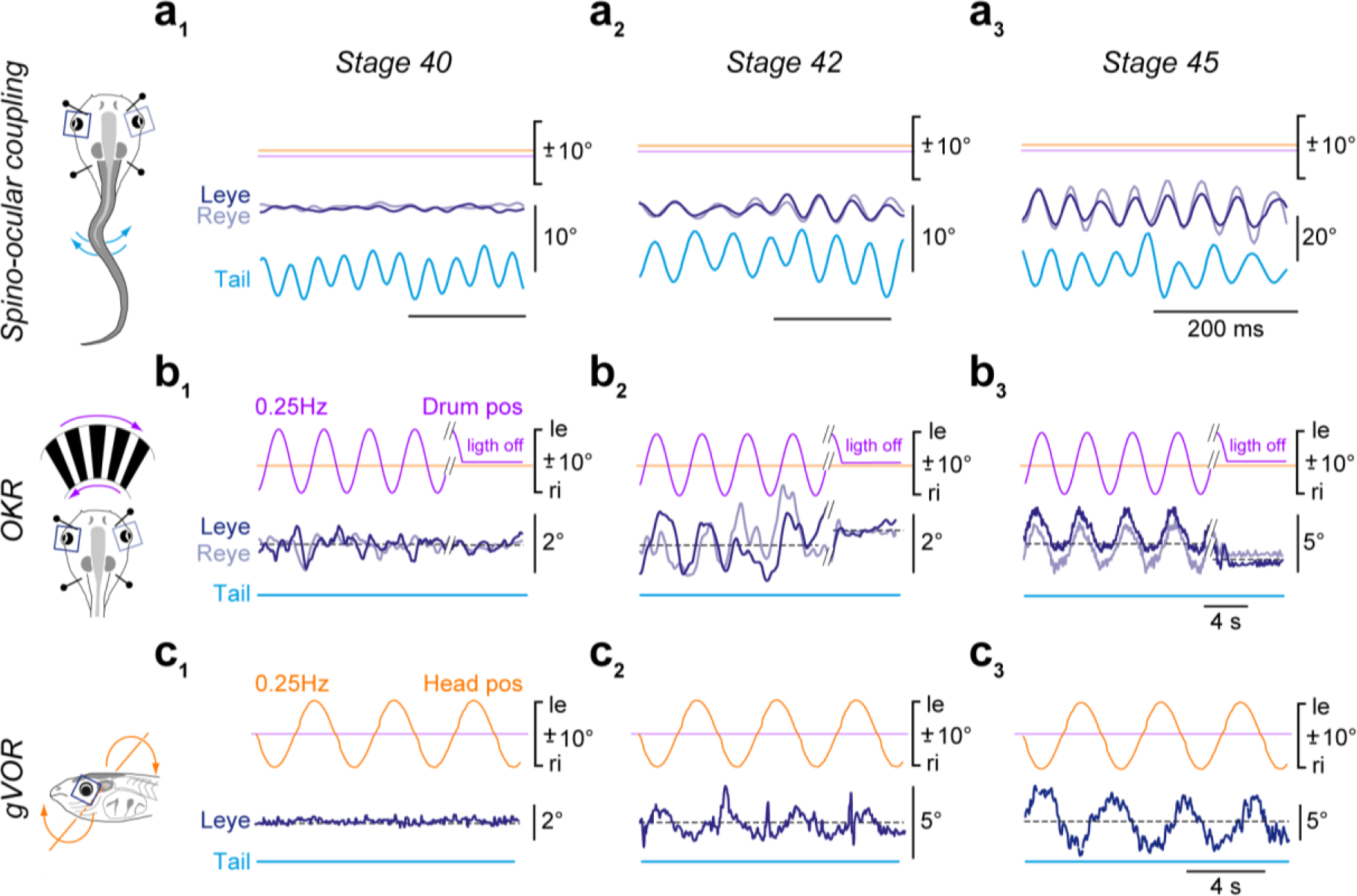
Common ontogenetic onset of spino-ocular, visual motion- and otolith organ-driven eye movements. **(a-c)** Gaze-stabilizing movements of the left (Leye, dark blue) and right eye (Reye, light blue) during spino-ocular motor coupling (a), optokinetic reflex (OKR, b) and gravitoinertial vestibulo-ocular reflex (gVOR, c) at stage 40 (a_1_, b_1_, c_1_), stage 42 (a_2_, b_2_, c_2_) and stage 45 (a_3_, b_3_, c_3_). Swim episodes occurred spontaneously; OKR and gVOR were elicited by sinusoidal rotation of black/white vertical stripes or forward-backward head roll motion at 0.25 Hz with positional excursions (pos) of ±10°. le, ri, leftward, rightward movement.

### Interactive maturation of locomotor-induced ocular motor behavior and angular vestibulo-ocular reflexes

Spino-ocular motor coupling experiences extensive spatio-temporal plasticity during amphibian metamorphosis, culminating at the emergence of appendicular and concurrent loss of axial muscle-based locomotor strategy in post-metamorphic froglets^10, 24^. To avoid interference with plasticity processes related to this latter final transition at stage 59, evaluation of the developmental plasticity in spino-ocular motor coupling was therefore restricted to larvae prior to this stage (Fig. 2a). At stage 49, ascending spinal locomotor efference copies, in the absence of motion-related visuo-vestibular sensory inputs, elicited phase-coupled oscillatory eye movements with rather invariable magnitudes of 5-7° (Fig. 2a_1_-b_1_) independent of tail bending amplitudes (*R* = 0.05 ± 0.05, *r*^*2*^ = 0.02 ± 0.02, *N* = 8; Fig. 2b_1_-b_2_). Consequently, the gain for locomotor efference copy-induced eye movements was relatively high for small tail excursions (0.74 ± 0.10 at left-right tail deflections of ∼10°, *N* = 8; Fig. 2b_3_, left panel) but decreased gradually for larger tail oscillations (Fig. 2b_3_; Supplemental fig. 2a). In contrast, at stage 58, eye and tail movement amplitudes were more closely correlated (*R* = 0.56 ± 0.11; *r*^*2*^ = 0.46 ± 0.07, *N* = 8; Fig. 2b_1_-b_2_) with a rather invariant gain of ∼0.4 independent of tail excursion magnitudes (Fig. 2b_3_, left panel; Supplemental fig. 2a). The spino-ocular motor coupling revealed an average phase lag *re* deflection of the rostral tail region of 0-30° (Fig. 2b_3_, right panel) that was unrelated to tail excursion magnitude as well as age (Watson-Williams F-test, *F* = 0.952, *p* = 0.337). The relative timing of the responses was rather variable at stage 49 but became considerably more homogeneous in older larvae (Supplemental fig. 2d,e, compare vector length of mean phase).

**Figure 2:**
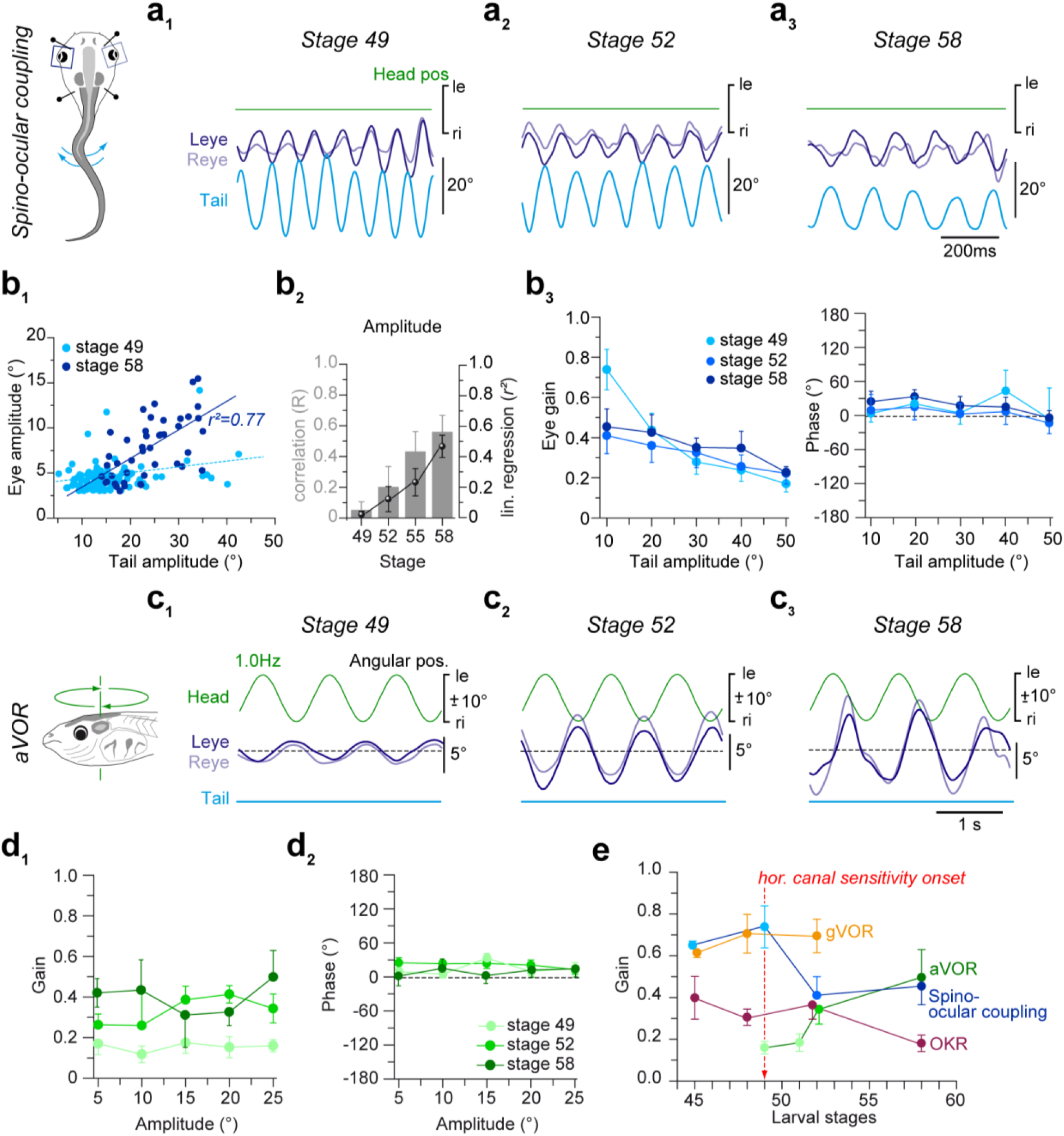
Developmental improvement of swim-related predictive ocular motor behavior and horizontal angular vestibulo-ocular reflex performance. **(a)** Movement of the left (Leye, dark blue) and right eye (Reye, light blue), elicited by spino-ocular motor coupling (pictogram) at stage 49 (**a**_**1**_), 52 (**a**_**2**_) and 58 (**a**_**3**_) during undulatory tail movements in head fixed preparations. **(b)** Quantification of spino-ocular motor coupling parameters presented as scatter plot of eye motion amplitudes as function of tail motion amplitudes (**b**_**1**_), histogram (grey bars) of the correlation coefficient (*R*) and linear (lin.) regressions (*r*^*2*^, black circles) between stage 49 and 58 (**b**_**2**_), linearity analysis of eye position gain (left in **b**_**3**_) and phase (right in **b**_**3**_) as function of tail undulation amplitude for three representative stages (see color-code). **(c)** Angular vestibulo-ocular reflex (aVOR) in response to sinusoidal horizontal (hor) head rotation (pictogram) at 1 Hz and ± 10° positional excursions at stage 49 (**c**_**1**_), 52 (**c**_**2**_) and 58 (**c**_**3**_) in the absence of swim-related tail undulations (light blue line). **(d)** Linearity analysis of the aVOR gain (d_1_) and phase (d_2_) as a function of head motion amplitude for three representative stages (see color-code) during sinusoidal rotations at 1 Hz. **(e)** Summary plot of eye position gain as index for the proficiency of spino-ocular motor coupling, aVOR, OKR and gVOR during larval development from stage 45 to 58. le, ri, leftward, rightward movement; all values represent mean ± SEM.

The developmental period between stage 49 and 58 was characterized by a considerable improvement of the performance of the horizontal angular vestibulo-ocular reflex (aVOR; Fig. 2c-d). First noticeable compensatory eye movements during horizontal head/body rotation were observed as early as stage 49-50 (Fig. 2c_1;_ *N=10*), although with a gain below 0.2, independent of stimulus strength (Fig. 2d_1,2_). This relatively late aVOR onset, compared to the gVOR and OKR is related to the necessity of semicircular canals to acquire sufficiently large duct diameters to become operational, which in *Xenopus* occurs shortly before stage 49-50^20^. Thereafter, aVOR gains became significantly increased by stage 52 (*N=6*), especially during larger amplitude head rotations (1 Hz, ± 15-25°; Fig. 2d,e) with further enhancement, although at slower rate, until stage 58 (Fig. 2d,e; *N=5*). The phase relation between semicircular canal-induced eye movements and stimulus waveform was relatively stable between stage 49 and stage 58, with constant small phase leads of the response *re* stimulus (∼0-30°) at 1Hz (Fig. 2d_1-2_).

Collectively, these findings suggest that implementation and refinement of locomotor efference copy-driven eye movements are closely intertwined with the developmental maturation of visuo- and vestibulo-ocular reflexes. This suggests that spino-ocular motor coupling co-emerges together with visuo- and otolith organ-driven ocular motor responses at stage 42 followed by progressive improvement in performance with a common time course until stage 49 (Fig. 1; Supplemental fig. 1). Thereafter, the further enhancement of locomotor efference copy-driven eye movements coincides with the maturation of the horizontal aVOR, with potentially cross-functional entanglement of the two ocular motor behaviors (Fig. 2e).

The potential impact of gradually improving aVOR performance on the maturation of locomotor efference copy-driven eye movements was elucidated in semicircular canal-deficient stage 52 larvae. These animals were obtained following bilateral injection of hyaluronidase into both otic capsules at stage 44 prior to inner ear duct completion^26^. This experimental perturbation at early larval stages caused a failure to form semicircular canals on both sides and consequently disabled the aVOR^25^. In compliance with the absence of functional semicircular canals, imposed horizontal head rotation at stage 52 failed to elicit compensatory eye movements (Fig. 3a), while in contrast locomotor efference copy-driven eye movements during tail undulations persisted (Fig. 3b). However, the performance of these predictive eye movements was inferior to that of age-matched controls and more comparable to that of younger larvae (Fig. 3c,d). The apparent retardation in spino-ocular motor performance might have derived from altered tail movements in semicircular canal-deficient stage 52 larvae, which were characterized by rather small amplitudes and high oscillation frequencies (Fig. 3e,f) reminiscent of those present in younger, stage 48 controls prior to aVOR onset (Fig. 3e,f)^30^. This suggests that semicircular canal signals are not only required for a functional aVOR, but also for the maturation of locomotor efference copy-driven predictive eye movements with stage-specific dynamic profiles, likely through a direct impact on locomotor pattern formation.

**Figure 3:**
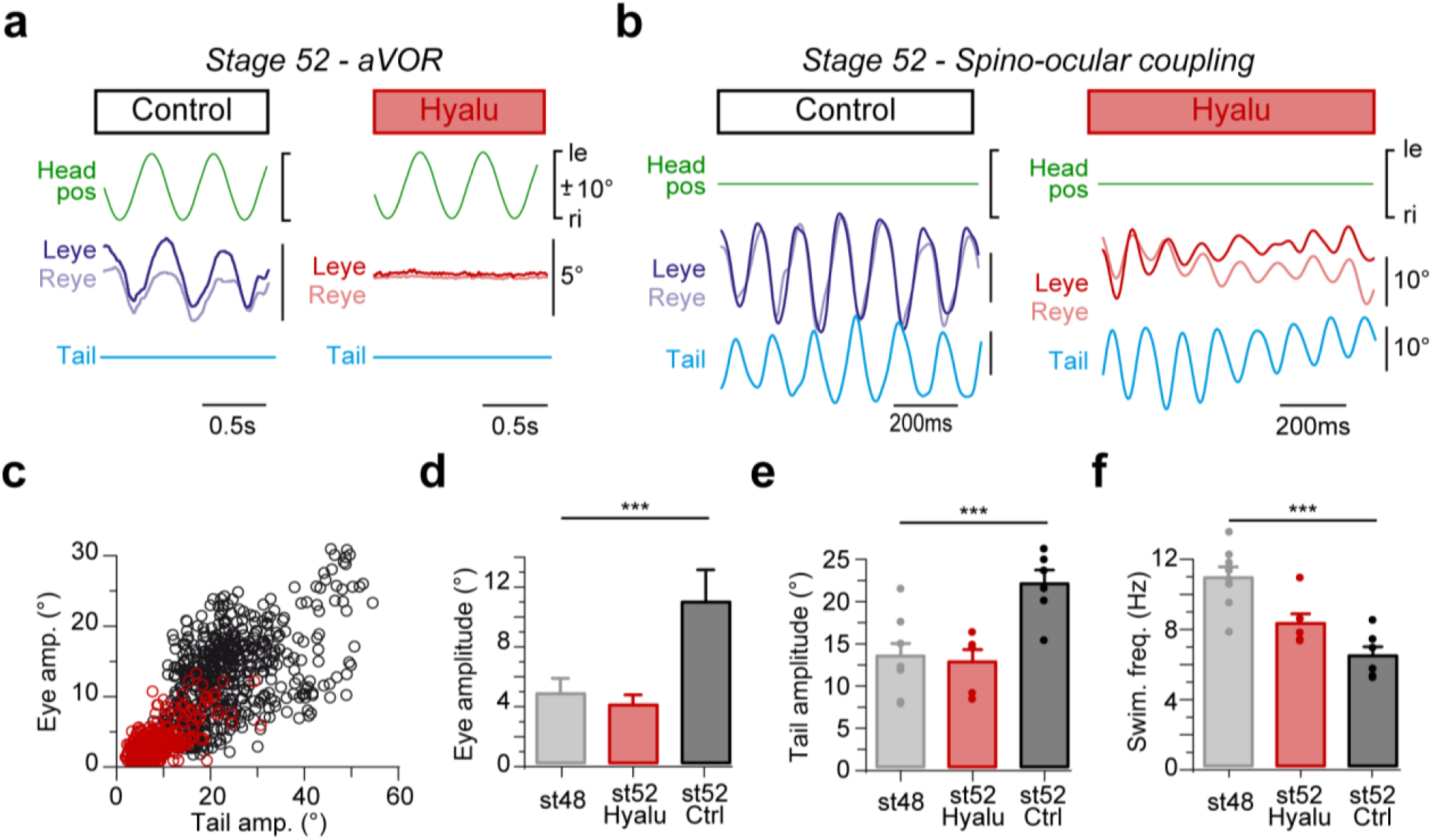
Semicircular canal deficiency prevents maturation of spino-ocular motor coupling. **(a)** Movements of the left (Leye, dark blue, red) and right eye (Reye, light blue, red) indicative for the angular vestibulo-ocular reflex (aVOR) evoked by horizontal sinusoidal head rotation at 1 Hz and ±10° positional (pos) excursion in the absence of swimming (**a**) and for episodes of spino-ocular motor coupling during tail undulations in a stationary preparation (**b**); recordings were obtained in a stage 52 control (left column in **a**,**b**) and an age-matched semicircular canal-deficient tadpole (right column in **a**,**b**), respectively; semicircular canal formation was prevented by bilateral intra-otic injections of hyaluronidase (Hyalu) at stage 44. **(c)** Quantification of locomotion-induced ocular motor coupling depicted as scatterplot of eye position amplitude as function of tail amplitude in stage 52 controls (black circles, *n=560 cycles*) and in hyaluronidase-treated age-matched animals (red circles, *n=375 cycles*). **(d-f)** Histogram of average eye position amplitude (**d**), swim-related tail undulation amplitude (**e**) and frequency (**f**) during spino-ocular motor coupling in stage 48 (light grey bars, *N=8*) and stage 52 controls (dark grey bars, *N=7*) and in hyaluronidase-treated stage 52 animals (red bars, *N=6*); statistical significance using Dunn’s multiple comparisons test, *** *p* < 0.05 is indicated; ns, not significant); le, ri, leftward, rightward movement.

### Progressive developmental influence of semicircular canals on locomotion-induced eye movements

The interaction between semicircular canal- and locomotor efference copy-driven eye movements was further evaluated by calculating the impact of horizontal head rotations on concurrent spino-ocular motor performance during undulatory swimming between developmental stages 45 and 58. Discrimination of vestibular and locomotor efference copy-driven response components in resultant eye movements was achieved by spectral analyses, which, however, required activation of the two components at different frequencies, respectively. Accordingly, passive head rotation was applied at 1 Hz (± 10° positional excursions), known to elicit an aVOR in larvae older than stage 49^20^. This stimulus frequency was sufficiently distant from swim-related tail beat oscillations that occurred within a range of 6-15 Hz^30^.

At stage 48, oscillatory eye movements during swim-related tail undulations and concurrent horizontal sinusoidal head rotation were robust (Fig. 4a_1_). However, noticeable 1 Hz vestibular motion stimulus-related eye movement components were absent as indicated by the lack of the respective spectral component (see absence of red band at 1 Hz in the spectrogram in Fig. 4b_1_ and of peak at 1Hz in the periodogram in Fig. 4c_1_). This lack is in compliance with the known failure to elicit an aVOR at this developmental stage^20^. The eye movement frequency component at stage 48 therefore corresponds exclusively to the swim frequency (∼12 Hz), indicated by the red band in the spectrogram and by the respective peak in the periodogram (Fig. 4b_1_,c_1_). As expected, at stage 52, eye movements during tail undulations and sinusoidal horizontal head rotation were also rather robust (Fig. 4a_2_). Spectral analysis demonstrated, however, that eye movements contained two clearly discernable components that coincided with the vestibular stimulus frequency (1 Hz) and the tail beat oscillations (∼6 Hz), indicated in the typical example by red bands in the spectrogram (Fig. 4b_2_) and corresponding peaks in the periodogram (Fig. 4c_2_).

**Figure 4:**
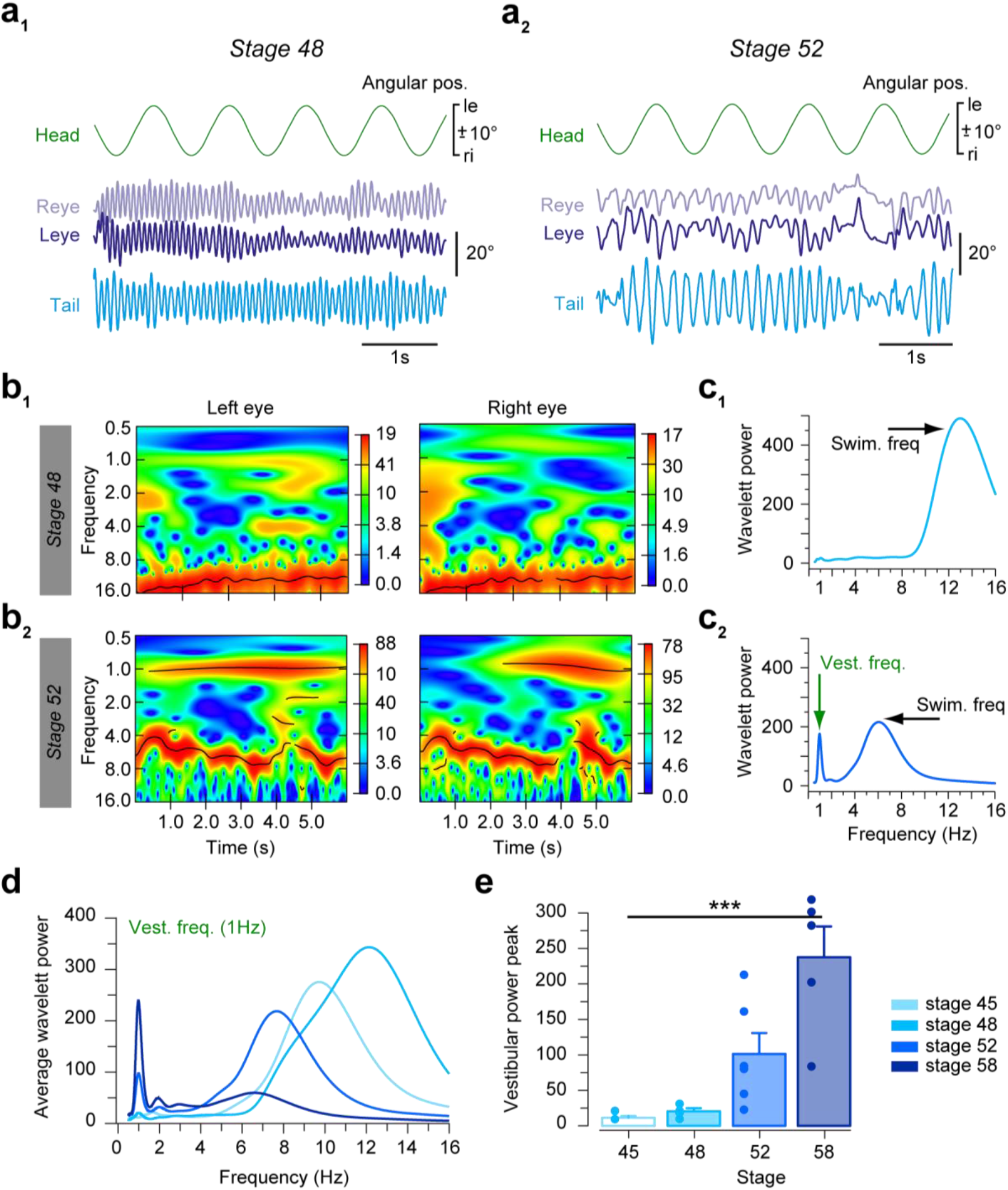
Progressive alteration of semicircular canal influence on gaze-stabilizing spino-ocular motor coupling during locomotion. (**a, b**) Representative recordings (**a**) and spectrogram (**b**) depicting the power spectrum of movement frequencies of the left (Leye, dark blue) and right eye (Reye, light blue) in head fixed preparations at stage 48 (**a**_**1**_, **b**_**1**_) and 52 (**a**_**2**_, **b**_**2**_) during swim-related tail undulations and concurrent horizontal head rotation at 1 Hz with positional excursions ± 10°; the occurrence of each frequency within the eye motion profile from 0.5 - 16 Hz is indicated by the color gradient from blue (rare) to red (often); black lines refer to significantly different wavelet power values within the spectrum. (**c**) Periodogram of two representative recordings at stage 48 (**c**_**1**_) and 52 (**c**_**2**_); note that eye movements at stage 48 consist of a single principal frequency that coincided with the tail undulations (Swim. freq. ∼13 Hz), while those at stage 52 contain two discernible frequencies consisting of tail undulations (Swim. freq. ∼6 Hz) and horizontal rotation (Vest. freq. 1 Hz). **(d, e)** Average periodogram of eye movements (**d**) during combined swim-related tail undulations and head rotation at 1 Hz at selected larval stages (color-coded); note the developmental increase of the frequency component related to the vestibular stimulus at 1 Hz, summarized in the histogram of average wavelet power amplitudes at stage 45 (*N=2*), 48 (*N=4*), 52 (*N=6*) and 58 (*N=5*) (**e**); all values represent mean ± SEM; number of animals is indicated in parentheses; le, ri, leftward, rightward movement.

Respective spectral analyses of eye movements during swimming and concurrent head rotation in larvae between stage 45 and 58 revealed a progressive reconfiguration of sensory-motor and predictive spino-ocular response components (Fig. 4d,e). Horizontal rotation-induced eye motion components experienced a gradual, yet significantly increasing contribution (Kruskal-Wallis test, *p* < 0.001, *N* = 17) after the onset of semicircular canal function at stage 49, culminating at stage 58, just prior to the metamorphic switch from tail-to limb-based locomotion (Fig. 4d,e). Possible underlying plasticity mechanisms include an increasing efficacy of peripheral and central processing of semicircular canal signals. This has a direct impact on extraocular motoneuronal activity as well as on spinal central pattern generator performance. A possible mechanism likely includes fusion of vestibular and locomotor efference copy signals at the spinal level before being transmitted by ascending pathways as joint premotor signal to extraocular motoneurons.

The superposition of vestibular and spino-ocular response components for the generation of eye movements during episodes of tail undulations and passive head motion was further specified in extraocular motoneurons as a more sensitive indicator (Fig. 5). The spike discharge of the left *lateral rectus* (LLR) nerve along with the motion of the right eye was recorded in semi-intact preparations at stage 52 (Fig. 5a-c). Fictive swimming, identified by episodes of rhythmic spike bursts in the right spinal ventral root (RVr), caused a phase-coupled spike discharge in the LLR nerve (Fig. 5a,b). Following integration at the neuromuscular interface of the eye muscles, the dynamics of this extraocular motor command corresponded to the horizontal oscillations of the intact right eye (Reye in Fig. 5b). Spectral analyses of the Reye movements and LLR nerve discharge revealed a synchronous oscillation frequency at ∼8 Hz, phase-coupled to the rhythm of the fictive swimming (Fig. 5c-e).

**Figure 5:**
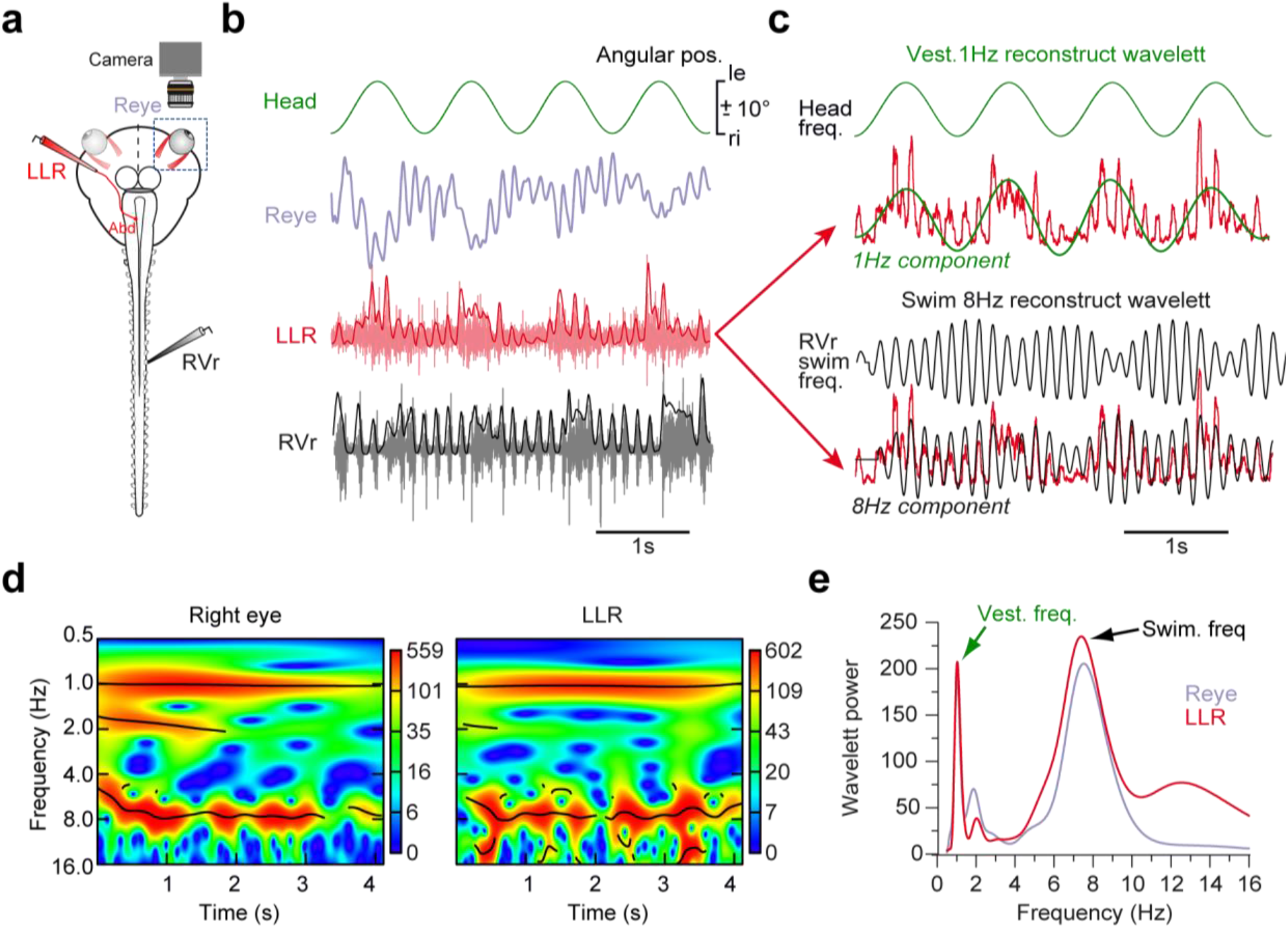
Eye movements and lateral rectus motor nerve discharge during vestibular stimulation and concurrent fictive swimming. **(a)** Simultaneous recording of ocular movements of the right eye (Reye), along with the spike discharge of the left lateral rectus (LLR, red) nerve and the right spinal ventral root (RVr, black) in a stage 52 preparation. (**b**) Representative recordings of eye movements LLR and RVr discharge during fictive swimming and horizontal head rotation (1 Hz, ± 10°) as spike activity (light red and grey) and after signal integration (dark red, black). **(c)** Wavelet reconstruction of vestibular stimulus-(top panel) and fictive swimming-related (bottom panel) frequency components extracted from the integrated LLR spike discharge after spectral analysis; the 1 Hz component (green trace superimposed on the red LLR discharge integral) matches head rotation (top panel) whereas the 8 Hz frequency component (black trace superimposed on red LLR discharge integral) coincides with the burst frequency of the fictive swim-related RVr spike discharge (bottom panel). **(d, e)** Spectrogram (**d**) and periodogram (**e**) depicting the vestibular motion stimulus-related 1 Hz (Vest. freq.) and the fictive swim-related 8 Hz frequency (Swim. freq.), expressed in the motion of the Reye and the spike discharge of the LLR nerve in **b** and **c**. Abd, abducens motor nerve; le, ri, leftward, rightward movement.

Swim-related LLR nerve spiking and eye oscillations were superimposed on a second, slower rhythm with a frequency of 1 Hz (Fig. 5c-e). This eye motion component corresponded to the head rotation-induced generation of ocular motor commands for an aVOR. Integration of signals related to swimming and concurrent sinusoidal head rotation, each with a different frequency, produced broadly tuned ocular motor oscillations with an amplitude modulation that matched the vestibular stimulus rhythm (Fig. 5c). The presence of two distinct response components (red bands in Fig. 5d and peaks in Fig. 5e) are thus of semicircular canal and spinal locomotor efference copy origin, respectively (Fig. 5e) and after fusion, jointly produce compensatory eye movements. Most interestingly, however, the cyclic Vr discharge was also broadly tuned, producing a spindle-shaped envelope with the timing of the head rotation frequency. This provides suggestive evidence that horizontal semicircular canal signals modulate the activity of spinal CPG circuits with direct implications for locomotor commands and ascending copies thereof, designated as ocular premotor signals.

### Developmental plasticity of vestibular influences on locomotor efference copy-evoked eye movements

Eye movements during swim episodes and simultaneous horizontal head rotation appeared to be governed by cross-modal interactions that principally fell into two categories (Figs. 6a,b, 7a,b). In the first category, typically found in younger larvae, sinusoidal head rotation during tail undulations were found to modulate the amplitude of individual swim cycle-related eye oscillations (Figs. 6a, 7a) to become alternatingly enhanced and attenuated depending on the head rotation half cycle (Fig. 7e, light blue dots). The variations (Δ amplitude, Fig.7i) were in the range of ∼2 - 8° between the largest and smallest eye excursions (5.14 ± 0.9°, *N=10*, Fig. 7i). In contrast, the position around which swim-related eye oscillations were centered remained relatively stable (Figs. 6a, 7a,f, light blue dots) with only small baseline positional fluctuations (Δ position, 2.34 ± 0.28°,*N=9*, Fig. 7j). The absence of larger, head rotation-timed eye oscillations suggests cancellation of these sensory signals, in compliance with earlier findings^8^. This outcome was further substantiated by recordings of extraocular motor activity from the LLR nerve (Figs. 6c, 7c). Individual efference copy-triggered spike burst amplitudes were modulated in phase-time with the head motion (Fig. 6c, reconstructed 1 Hz wavelet; 7c; light red dots in Fig. 7g; Δ amplitude, 178.7 ± 120.2%, *N=5*, Fig. 7k). In contrast, the baseline spike rate remained rather unaffected by the head rotation, illustrated in the typical example as minor vestibular stimulus-timed fluctuations of the discharge across a single motion cycle (Fig. 7c; light red dots shown in Fig. 7h; Δ baseline, 77 ± 20%, *N=5*, Fig. 7l).

**Figure 6:**
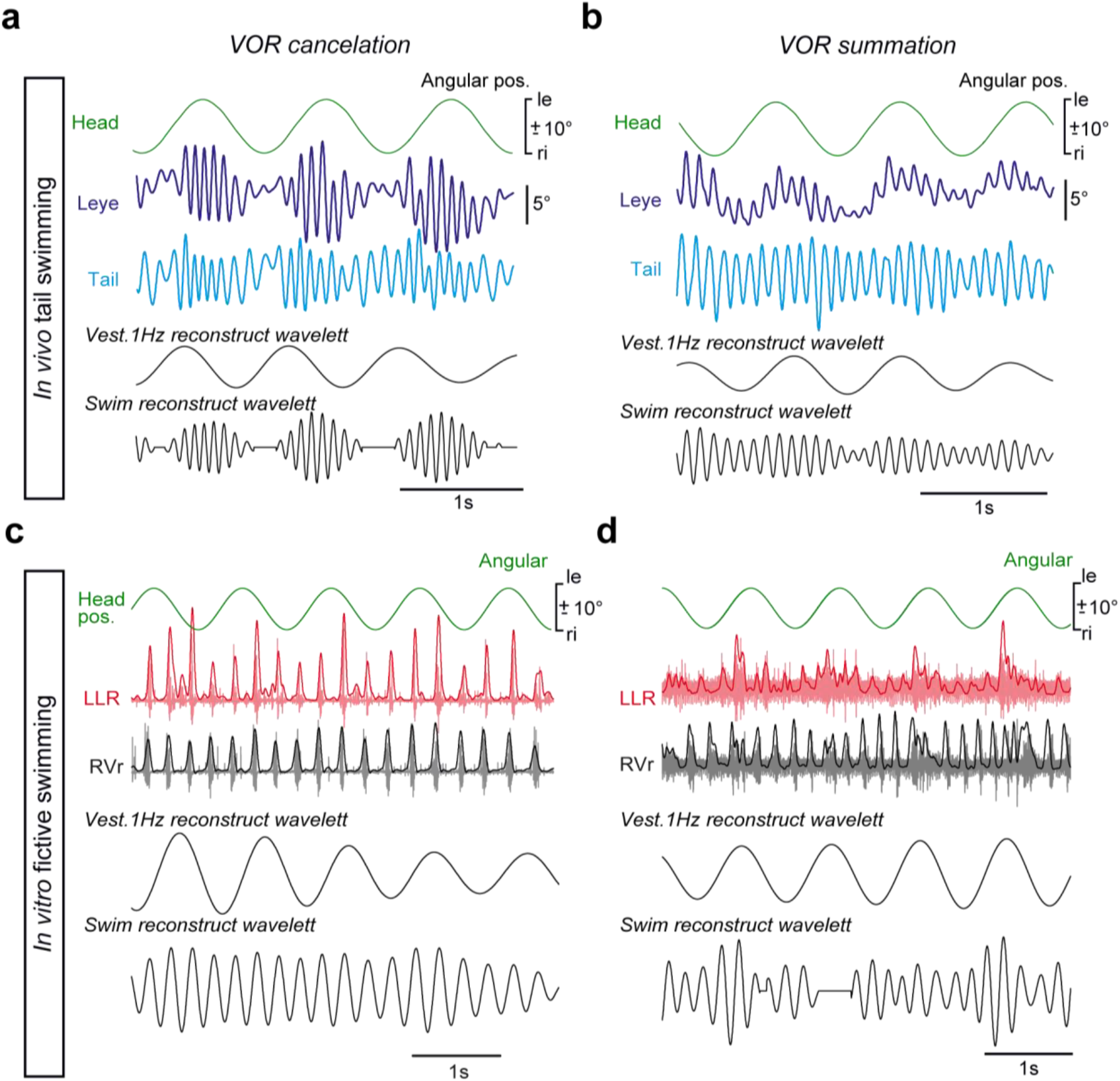
Developmental plasticity of vestibular and spino-ocular motor signal integration during gaze-stabilization. (**a, b**) Representative examples of movements of the left eye (Leye) during horizontal head rotation and concurrent tail undulations, depicting either a cancellation of the angular vestibulo-ocular reflex (aVOR; **a**) by spino-ocular motor signals or summation of both eye motion components (**b**); black traces represent wavelet reconstructions of the vestibular stimulus and the swim-related component, extracted from the Leye motion recording by spectral analyses, respectively. **(c, d)** Representative examples of spike discharge recorded from the left lateral rectus (LLR) nerve during horizontal head rotation and concurrent fictive swimming, indicated by the rhythmic discharge of the right spinal ventral root (RVr); aVOR-related LLR nerve spike discharge was either canceled by (**c**) or summated (**d**) with spinal locomotor efference copies during fictive swimming; black traces represent wavelet reconstructions of the vestibular stimulus and the fictive swim-related component, extracted from the LLR nerve spike discharge by spectral analyses, respectively. le, ri, leftward, rightward movement.

**Figure 7:**
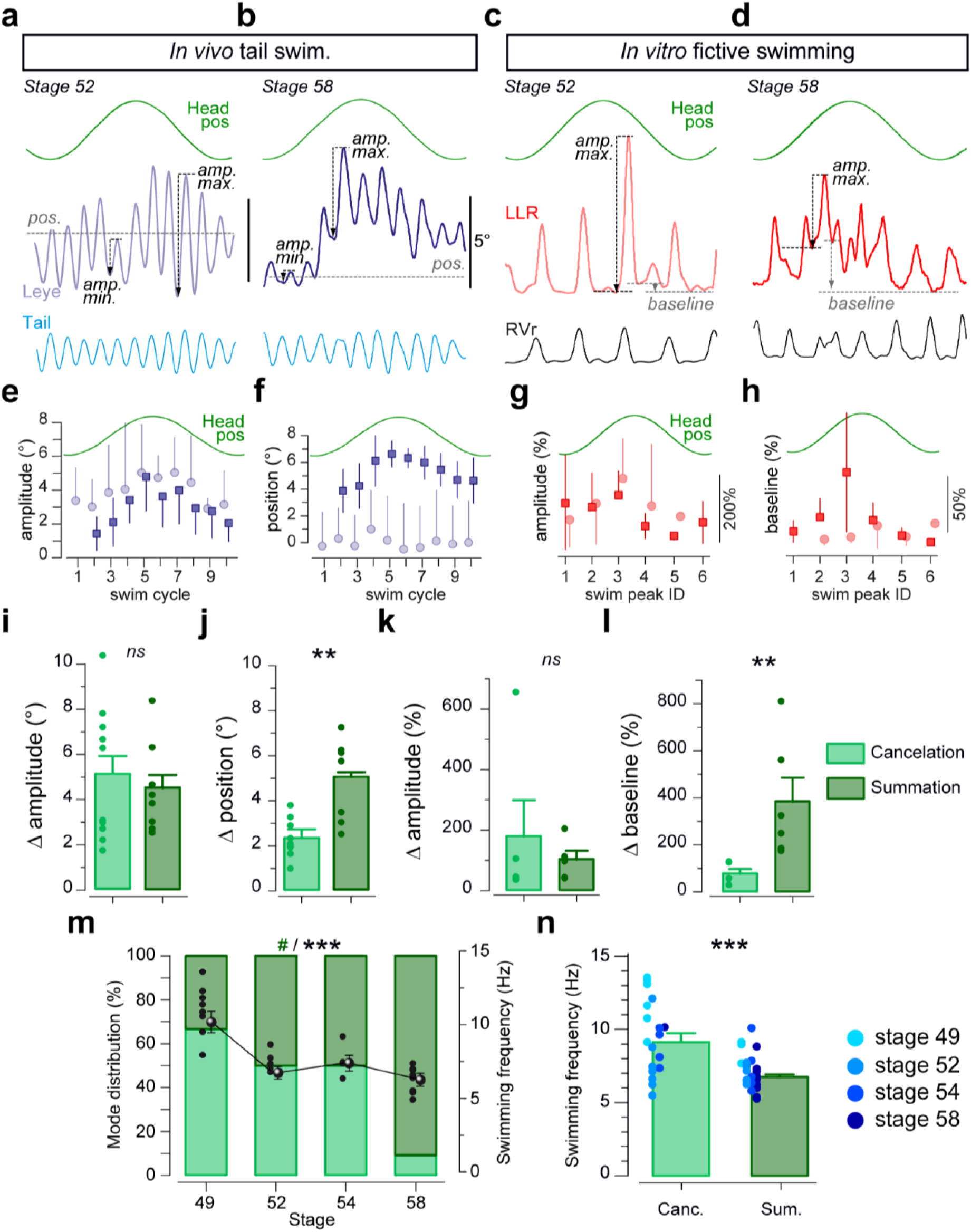
Computational characteristics of vestibulo-ocular and spino-ocular motor signal integration. **(a, b)** Movements of the left eye (Leye) illustrating spino-ocular motor coupling during tail undulation over a single horizontal head rotation cycle (Head pos) in a stage 52 and 58 tadpole; eye movements either result from a cancelation of vestibulo-ocular signals (**a**) or from a summation of vestibulo-ocular and spino-ocular signals (**b**) with differential effects on the amplitude (amp.) and baseline position (pos.). **(c, d)** Discharge of the left lateral rectus (LLR) nerve depicting spino-ocular motor coupling during rhythmic spinal ventral root (RVr) discharge (fictive swimming) over a single horizontal head rotation cycle (Head pos) in a stage 52 and 58 tadpole; cancelation of vestibulo-ocular signals (**c**) or summation of vestibulo-ocular and spino-ocular signals (**d**) can be dissociated by the differential impact on the discharge amplitude (amp.) and baseline position (pos.), respectively. (**e, f)** Average (± SD) of amplitude (**e**) and baseline position (**f**) of swim cycle-related Leye movements over a single head rotation cycle during vestibular cancelation (light blue dots, *n=6 cycles*) and summation (dark blue square, *n=6 cycles*), respectively. **(g, h)** Average (± SD) of burst amplitude (**g**) and discharge baseline position (**h**) of the RVr burst-related LLR discharge over a single head rotation cycle during vestibular cancelation (light red dots, *n=6 cycles*) and summation (dark red square, *n=6 cycles*), respectively. (**i, j**) Histogram of maximal ocular motion amplitude variations (Δ amplitude, **i**) and baseline position variations (Δ position, **j**) over a single vestibular motion cycle during swimming and concurrent head rotation. **(k, l)** Histogram of the integrated maximal LR nerve burst amplitude variations (Δ amplitude, **k**) and maximal baseline discharge variation (Δ baseline, **l**) over a single vestibular motion cycle during fictive swimming and concurrent head. **(m)** Relative occurrence of “cancelation” and “summation” (left axis, Chi-square test, #, *p* < 0.05) and average swimming frequency (right axis; stage 49 *N=9*, 52 *N=14*, 54 *N=9*, 58 *N=11*; Kruskal-Wallis test, ***, *p* < 0.001) between stage 49 and 58; number of animals is indicated in parentheses. (**n**) Histogram of the mean swim frequency calculated for 18 locomotion-induced ocular motion sequences expressing cancelation and 25 sequences expressing summation, color-coded with respect to larval stages.

Cross-modal interactions that fell into the second category were typically found in older larvae (Fig. 6b,d). Amplitudes of individual swim cycle-related eye oscillations and corresponding LLR nerve motor commands were modulated by the head rotation. The overall pattern was comparable to that of the first category (Fig. 7b,d), although with smaller amplitude variations between the largest and smallest eye oscillations (Fig. 7b; dark blue squares in Fig. 7e; Δ amplitude, 4.5 ± 0.7°, *N=8*, Fig. 7i). Most noticeable, however, the baseline eye position around which swim-related eye movements oscillated was considerably modulated with a waveform that corresponded to the head rotation, causing considerable left-right eye movements (Fig. 7b; dark blue squares in Fig. 7f; Δ position, 5.05 ± 0.62°, *N=8*, Fig. 7k). This eye movement profile complies with a summation of VOR components and locomotor efference copy-triggered eye oscillations during joint swimming and passive head rotation. This eye motion pattern matched corresponding variations of the spike discharge of the LLR nerve (Figs. 6d, 7d). Locomotor efference copy-triggered spike burst amplitudes as well as the baseline spike activity were modulated during the rotation with a profile that was phase-timed to the head motion (Fig. 7g,h, dark red squares; Δ amplitude, 102,3 ± 29.7%, *N=5*; Δ position, 382.6 ± 103.4%, *N=6*) as compatible with the eye oscillations (Fig. 7e,f, dark blue squares).

Quantification of the two patterns of signal integration principles across a larger population of animals revealed that the vestibular influence on efference copy-triggered eye oscillations and associated spike bursts were independent of larval age (Fig. 7i,k; Mann Whitney *U*-test, *p* = 0.910 and *p* = 0.690 respectively) and thus not indicative for either one of the two integrative principles. However, the vestibular influence on baseline eye position and extraocular motor spiking was significantly larger in the typical example obtained from older tadpoles (Fig. 7j,l; Mann Whitney *U*-test, *p* < 0.01). This difference represents a distinguishing feature of the cross-modal signal computation during swimming and motion stimulation (Fig. 6). In fact, extraocular motor signals cancellation *versus* summation were typically related to younger *versus* older tadpoles, respectively (Fig. 7m). Although, either one of the two modes could be found in tadpoles of any age (Sup. Fig. 6), signal cancellation (light green in Fig. 7m) predominated in younger tadpoles (∼66% of stage 49 larvae, Fig. 7m) but was rarely observed in older larvae (∼10% of stage 58 larvae, Fig. 7m). Conversely, additive signal interactions (summation; dark green in Fig. 7m) of vestibular and intrinsic locomotor-induced extraocular motor signals correspondingly predominated in older larvae (90% at stage 58, Fig. 7m).

Despite the clear evidence for an age-related shift in signal processing (light and dark green bars in Fig. 7m), larval age might only be an indirect classifier. As demonstrated earlier^9, 30^, the frequency of swim-related tail oscillations decreases progressively during larval development (black dots and bowls in Fig. 7m). Thus, using swim frequency as decisive parameter, cancellation of motion-related ocular motor response components by locomotor efference copies occurred mainly at higher tail undulatory frequencies (∼10 Hz, black dots and bowls in Fig. 7m), while summation of signals was found, whenever swimming occurred at a low frequency such as in stage 58 larvae (∼6 Hz, black dots and bowls in Fig. 7m). Accordingly, swim episodes in which vestibular signals were additively combined with locomotor efference copies had significantly lower frequencies (Mann Whitney *U*-test, *p* < 0.001; 6.77 ± 0.24 Hz, Fig. 7n) than those where concurrent vestibulo-ocular signals were canceled (9.16 ± 0.60 Hz, Fig. 7n), independent of developmental stage (see color-coded dots in Fig. 7n).

This differential interaction therefore provides suggestive evidence that the interaction between locomotor efference copies and vestibular sensory signals gradually alters during larval development. In young tadpoles at stage 49-50, horizontal semicircular canal signals modify spinal CPG activity, while the VOR-related motor output is suppressed. In older animals, at stage 58, VOR cancelation by locomotor efference copies disappears and instead is replaced by a summation of intrinsic and sensory-motor components in the process of generating extraocular motor commands. Key to this developmental switch is a gradually reduced efficacy or rhythmicity of the locomotor program, illustrated by the slower tail undulation during metamorphic progression with concurrent vanishing influences of locomotor efference copies on the ocular motor output.

## DISCUSSION

Locomotion-driven gaze-stabilizing eye movements appear concomitantly with visual- and otolith organ-driven ocular motor reflexes during early larval life in *Xenopus* but are tuned by a second step of maturation initiated by the acquisition of semicircular canal sensitivity. With the appearance of a robust angular VOR, rhythmic ocular movements driven by locomotor CPG efference copies experience a gradual alteration in the modulatory influence of semicircular canal signals during larval developmental. During swimming episodes with high-frequency tail undulations, preferentially occurring in young larvae, horizontal head rotations are suppressed by locomotor efference copies, causing selective expression of ocular rotations around a rather invariant baseline eye position. Older larvae, which predominantly swim with low-frequency tail undulations, exhibit ocular motor reactions that result from a summation of horizontal head rotational signals and locomotor efference copies. The corresponding computational regime thereby produces gaze-stabilizing conjugate eye movements that are a combination between the angular VOR and spino-ocular motor coupling.

### Common developmental time course of sensory-based and locomotion-driven gaze stabilization

Larval *Xenopus* acquire the capacity to swim freely around stage 37/38 when spinal motor networks are sufficiently mature to allow the production of muscular activity that produces rostro-caudally progressing undulation of the tail (Fig. 8a; ^28, 31, 32, 33^). However, locomotion-driven eye movements were only observed at stage 42-43, despite the earlier onset of spinal CPG activity (Fig. 8a). Interestingly the onset of locomotion-driven ocular motor behavior concurs with the ontogeny of visual-driven ocular reflexes and otolith organ-dependent VOR components (Fig. 8a; ^19, 20^). While the vestibular sensory periphery continues to grow after hatching until late pro-metamorphic stages, the morphogenesis of the labyrinthine endorgans occurs early during ontogeny, concomitantly with central vestibulo-motor connections^34, 35^. Based on data from chicken embryos, the first synapse to form along the VOR three-neurons arc appears to be the connection between extraocular motoneurons and eye muscles^36^. Subsequently, vestibular afferent fibers connect to central vestibular neurons before the latter neurons form synaptic connections with extraocular motoneurons. This sequence complies with the onset of otolith organ-dependent reflexive eye movements. Hence, from stage 42 onward, the maturation of otolith organs allows detection of gravitoinertial head movements^15, 20^. In contrast, vestibular projections to the spinal cord are fully established at stage 42-43^37^. Thus, the occurrence of first locomotion-driven compensatory eye movements could depend on established otolithic VOR central connections as well as the functionality of the respective endorgan. If this logic reflects the full picture is unknown so far. In fact the developmental switch from a single spike motor output to a burst pattern between stage 37/38 and stage 42 could supplement the increased spino-ocular motor coupling during this period (Supplemental Fig. 1a; ^33^).

**Figure 8:**
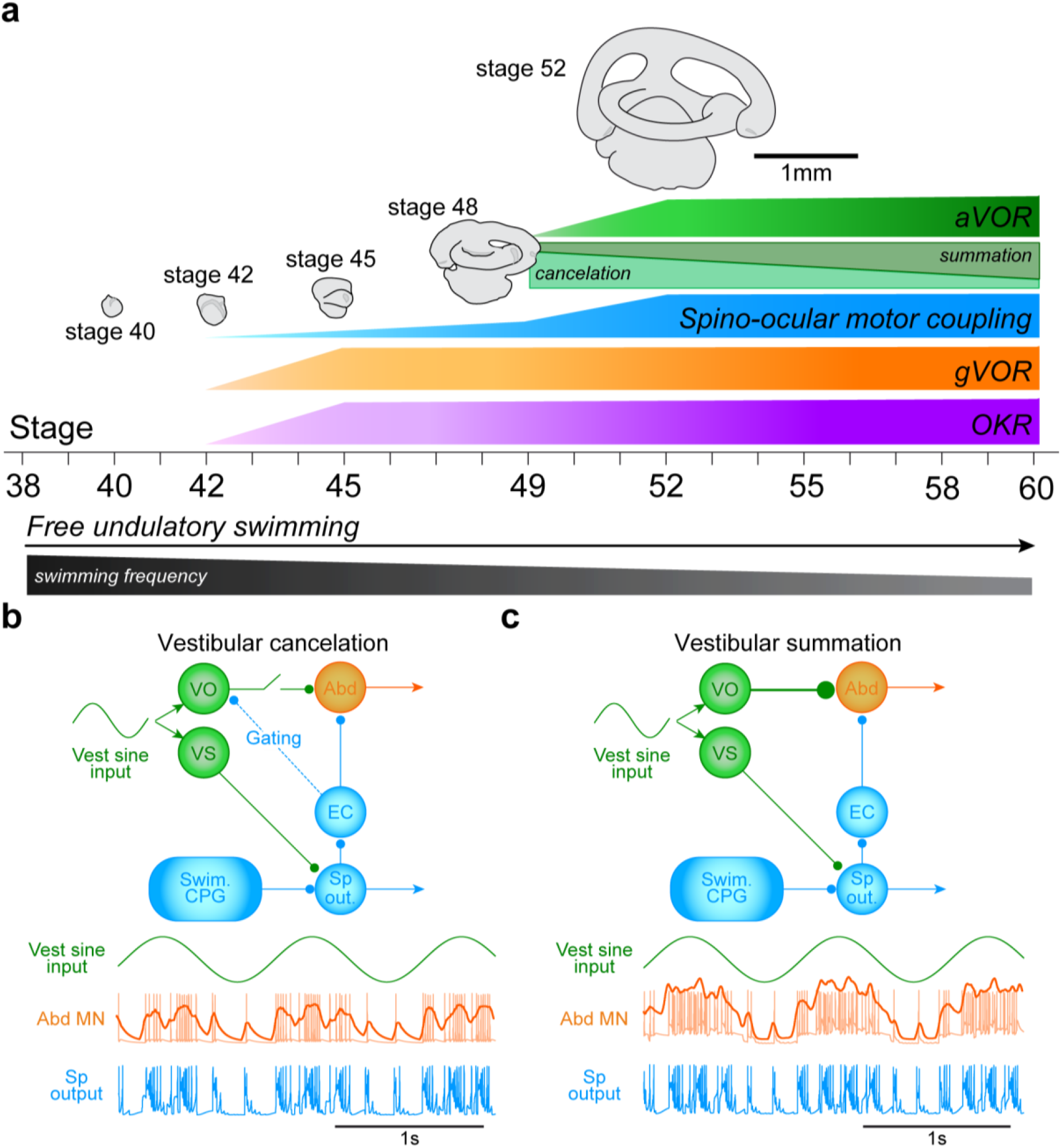
Developmental plasticity of locomotion-related spino-ocular and visuo-vestibulo-ocular motor computations. (**a**) Summary scheme depicting the time frame of the ontogenetic maturation of angular and gravitoinertial vestibulo-ocular reflexes (aVOR, gVOR), the optokinetic reflex (OKR and swim-related spino-ocular motor coupling across larval stages and growth of inner ear endorgans. (**b, c**) Neuronal activity of abducens motoneurons (Abd MN) and spinal motor output (Sp out.), constructed by a NEURON network model based on putatively involved brainstem and spinal circuit elements; the model robustly differentially reproduced the cancelation of vestibulo-ocular signals (**b**) as well as the summation of vestibulo-ocular and spino-ocular signals (**c**) during swimming, depending on swim rhythm magnitudes. CPG, swim central pattern generator; EC, efference copy neuron; Vest sine input, vestibular sinusoidal input; VO, vestibulo-ocular; VS, vestibulospinal.

### Spino-ocular motor function depends on a joint developmental progression of otolith and semicircular canal organs

After the onset of the aVOR, spinal CPG-driven eye movements become more efficient in the compensation of tail undulatory movements at all frequencies providing appropriate amplitude and temporal relationships (constant gain and minimal phase shift; see Fig.2b and supplemental fig.2). The inappropriate amplitude and the phase shift of locomotion-induced eye movements in hyaluronidase-treated and thus semicircular canal-deficient larvae suggests that this second maturation step depends on the integrity of the ducts (see Fig. 3). Unlike semicircular canals related vestibulo-ocular pathways where the different ducts and eye muscles pairs share a coplanar spatial organization^38, 39^, macular VOR circuits have to transform an otolithic planar sensitivity with a 360° directional distribution onto spatially appropriate sets of extraocular motoneurons with directionally matching pulling directions ^40^. This major difference could explain the two steps in the maturation process of locomotion-induced ocular motor behaviors, before and after the emergence of the aVOR at stage 49/50. Therefore, given the absence of matching sensory and extraocular motor frameworks for otolith signals, semicircular canal signals could be the relevant sensory signal to tune efference copy-based ocular motor behaviors at later developmental stages (Fig. 8a). In fact, the spatial tuning of otolith signals in extraocular motoneurons depends on a co-activation of specific epithelial sectors of the utricle, spatially aligned with semicircular canals ^25^. As supportive evidence for this assumption, the spatial tuning of utricular signals coincides with the appearance of the aVOR and is abolished in semicircular canal-deficient *Xenopus* larvae ^25, 41^. This spatial refinement of otolith signals by semicircular canal signals could also assist in the observed semicircular canal-dependent second maturation step of locomotion-induced eye movements.

### Vestibular computations responsible for different modes of locomotion-driven ocular motor behaviors

The current data demonstrate that semicircular canal sensory signals were either cancelled by locomotor efference copies or integrated with spino-ocular motor-driven eye movements. Although both computational modes can be observed in a given animal, depending on the magnitude of the swim frequency (see Supplemental fig. 6), the expression of either cancelation or summation is largely related to the developmental stage. Cancellation of vestibular signals dominates at stage 49-51 when the aVOR is not yet fully established and the animals express high-frequency swimming, whereas a summation dominates at stage 58 when the aVOR is robust in animals that swim at lower frequencies (Fig. 6 and 7). These findings concur with previous observations demonstrating that the lateral rectus nerve discharge is temporally gated by the CPG efference copies during combined fictive swimming and horizontal head rotation at stage 52-53 ^8^.

The decision to cancel or summate signals obviously derives from a swim frequency-dependent differential gating of vestibular sensory inputs and locomotor efference copies. During high-frequency tail undulations, compensatory eye movements reflect entirely spino-ocular motor commands, while horizontal semicircular canal inputs appear to only adjust the amplitude of the ocular excursion. This restrained vestibular modulatory effect could be mediated by vestibulo-spinal inputs on spinal locomotor networks, generated by the semicircular canal activation during head rotations. During low-frequency tail undulations, horizontal semicircular canal inputs modulate both the spino-ocular motor signals as well as the baseline ocular position, likely due to a weaker gating of vestibular sensory signals by efference copies. This latter feedback-feedforward interaction of motion signals during swimming produces compound compensatory eye movements that are a mixture of aVOR and locomotion-induced ocular motor behavior. If the efference copy-driven gating of vestibular inputs is produced by an inhibitory action on the horizontal aVOR at the level of central vestibular neurons or directly at the level of extraocular motoneurons remains to be investigated. The fusion of spino-ocular and vestibulo-ocular signals could be the result of synaptic integration of VOR signals and those of the ascending spino-ocular pathway that both of which terminate directly on abducens motoneurons. We wanted to explore this hypothesis by using a computational neuron network modeling the principal connections between vestibulo-ocular, vestibulo-spinal and spino-ocular pathways (Fig. 8b, c). In the vestibular cancelation mode (Fig. 8b) the direct vestibular input to Abducens motoneuron was turn off to mimic the spinal gating of the aVOR during swimming^8^. The only remaining vestibular input was the vestibulospinal neuron connecting to the spinal output neuron. In this model configuration, the Abducens motoneuron exhibited a discharge pattern during combined fictive swimming and vestibular activation (1Hz) comparable to LR motor nerve discharge shown in figures 6c and 7c, with a weak modulation of the discharge baseline. Inversely, in the vestibular summation mode (Fig. 8c), direct vestibulo-ocular inputs were not turn off on Abducens motoneurons. As a consequence the imposed vestibular stimulation (1Hz) strongly modulated the discharge baseline of the Abducens motoneuron during fictive swimming activity, in a comparable manner that the LR motor nerve discharge recorded during vestibular-swimming summation situations presented in figures 6d and 7d. This restrained computational experiment, based on a simple modeling neuronal network, constitutes a promising preliminary result to support our hypothesis claiming that cancelation and summation interaction modes between vestibular and locomotor signals could be relayed by distinct brainstem-spinal pathways, without excluding the effect of other synaptic interactive effects.

The complex interaction of vestibular- and spinal CPG-driven compensatory eye movements is the likely cause for optimal gaze-stabilization also in terrestrial vertebrates including humans ^13^. In fact, the predominance of spino-ocular motor behavior during rhythmic walking appears to be a key feature of gaze-stabilization. However, in contrast to quadrupedal locomotion, the study of a simpler and more accessible model, such as larval *Xenopus* allows demonstrating the ontogenetic establishment, maturation and tuning of associated feedforward gaze-stabilizing motor function. In a comparable manner, the implication of the vestibular sense has recently been demonstrated for the development of locomotor coordination and postural control ^42, 43^. In summary, the current results generate a unique hypothesis about feedback-feedforward interactions between vestibular and locomotor signals for terrestrial animals with bi- and quadrupedal locomotor patterns such mice ^11^.

## ACKNOWLEDGEMENTS

This work was supported by grants from the *Centre National de la Recherche Scientifique* and by the *Agence National pour la Recherche* (Locogaze - ANR-15-CE32-0007-01). FML received support from the INCIA CNRS UMR5287. The authors are grateful to Lionel Para-Iglesias for his work in the animal facility and for his help in 3D printing (*Creativ* core facility), to Philippe Chauvet from the INCIA mechanical workshop and to Daniel Cattaert to help designing the computational network model with the NEURON software. The authors thank Dr. Romuald Nargeot for his help with the statistical analyses of the data and the development of spectral analysis tools.

## AUTHOR CONTRIBUTION

Designed research: FML - DC – HS

Performed research: FML - JBC – GC

Analyzed data: FML - JBC - GC - MB

Wrote the paper: FML - JBC - DC - HS - MB

## DATA AVAILABILITY

Further information and requests should be addressed to the corresponding author François M. Lambert.

---

**Supplemental figure 1:**
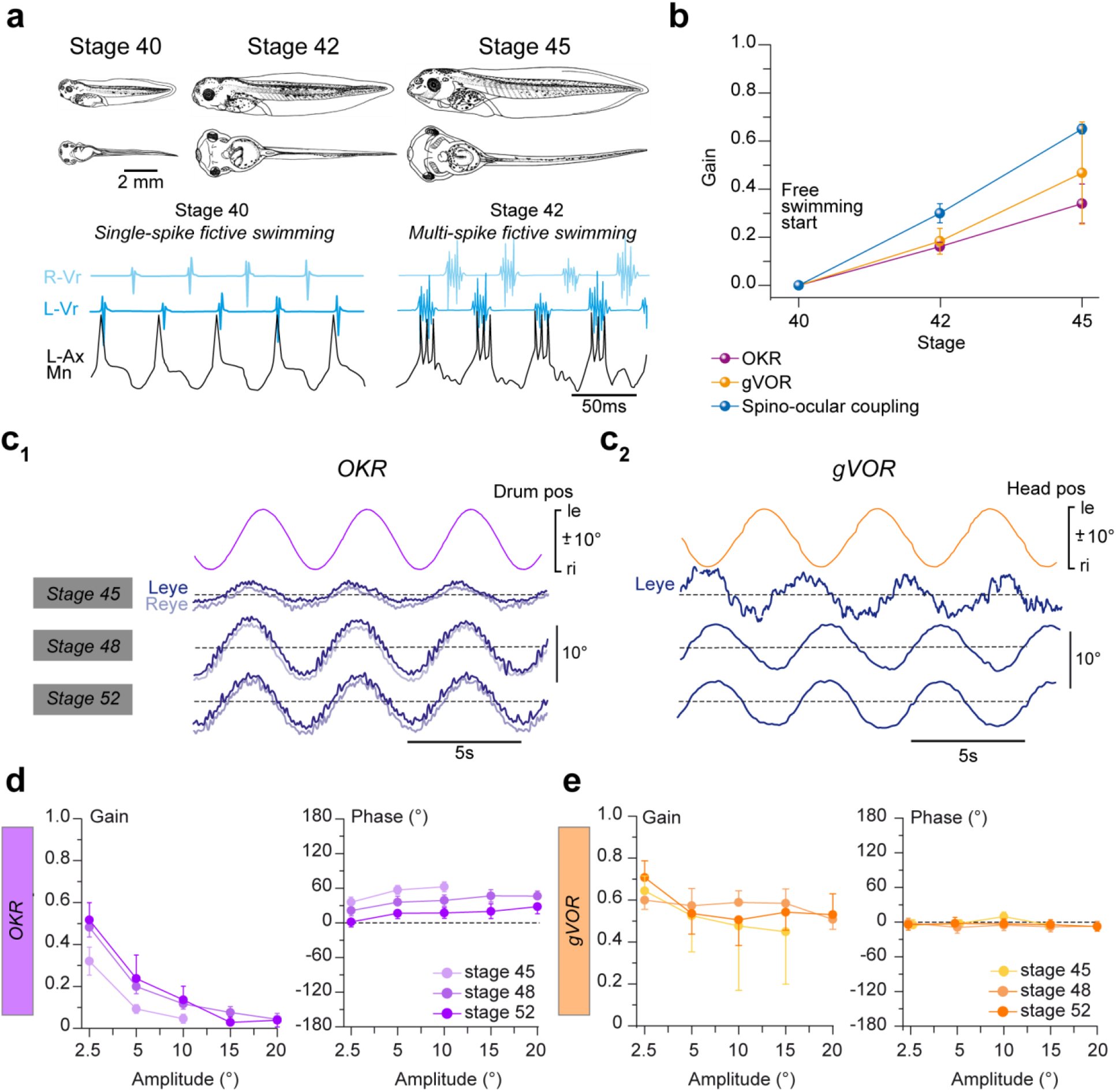
Ontogenetic improvements of gaze-stabilizing ocular motor behaviors. **(a)** Developmental switch from single-spike to multi-spike activity in spinal axial motoneurons (Ax Mn) between stage 40 and 42 (modified from ^29^). **(b)** Eye position (pos) gain for OKR, gVOR and spino-ocular motor coupling at stage 40, 42 and 45. **(c)** Eye movements of the left (Leye, dark blue) and right eye (Reye, light blue) during the optokinetic (OKR, **c**_**1**_) and gravitoinertial vestibulo-ocular reflex (gVOR, **c**_**2**_) at stage 45, stage 48 and stage 52; ocular motor responses were elicited by sinusoidal rotation of black/white vertical stripes or forward-backward head roll motion at 0.25 Hz with positional excursions (pos) of ± 10°. **(d**,**e)** Eye position gain (left) and phase (right) for the OKR (**d**) and gVOR (**e**) as function of the stimulus amplitude. le, ri, leftward, rightward movement; all values represent mean ± SEM.

**Supplementalfigure 2:**
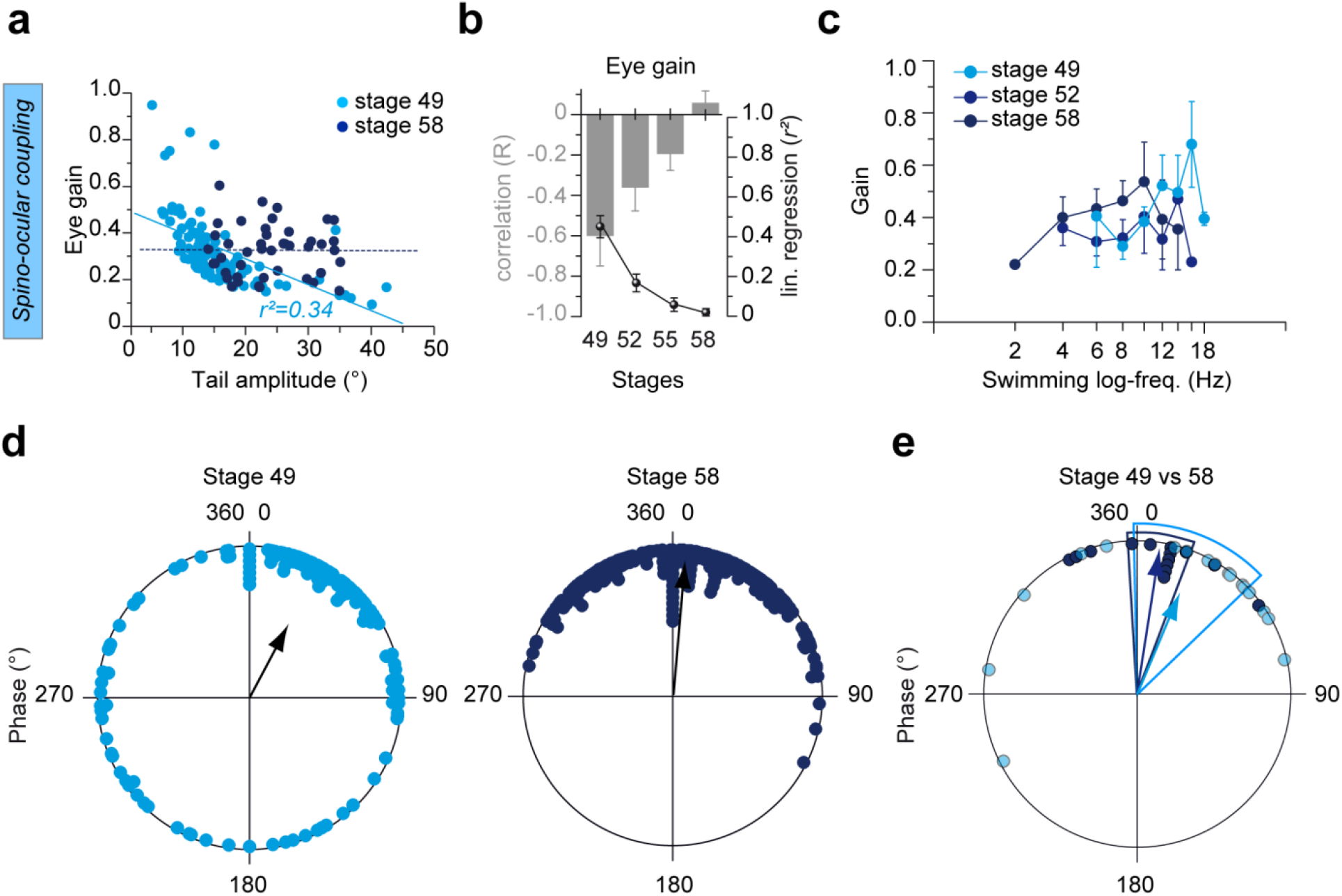
Developmental enhancement of spino-ocular motor coupling. **(a)** Scatterplot representing eye position gain as function of tail undulation amplitude at stage 49 and 58. **(b)** Histogram (grey bars) of correlation coefficients (*R*) and linear (lin.) regression (*r*^*2*^, black circles) for eye position gains at stage 49, 52, 55 and 58. **(c)** Bode analysis of eye position gain (mean ± SEM) as function of tail undulation frequency at stage 49, 52, 58. **(d)** Polar plot of phase relations between eye and tail positions for a representative example at stage 49 (left, light blue) and stage 58 (right, dark blue). **(e)** Polar plot of average (mean ± SEM) phase relations at stage 49 (*p* < 0.001) and stage 58 (*p* < 0.001). le, ri, leftward, rightward movement.

**Supplemental figure 6:**
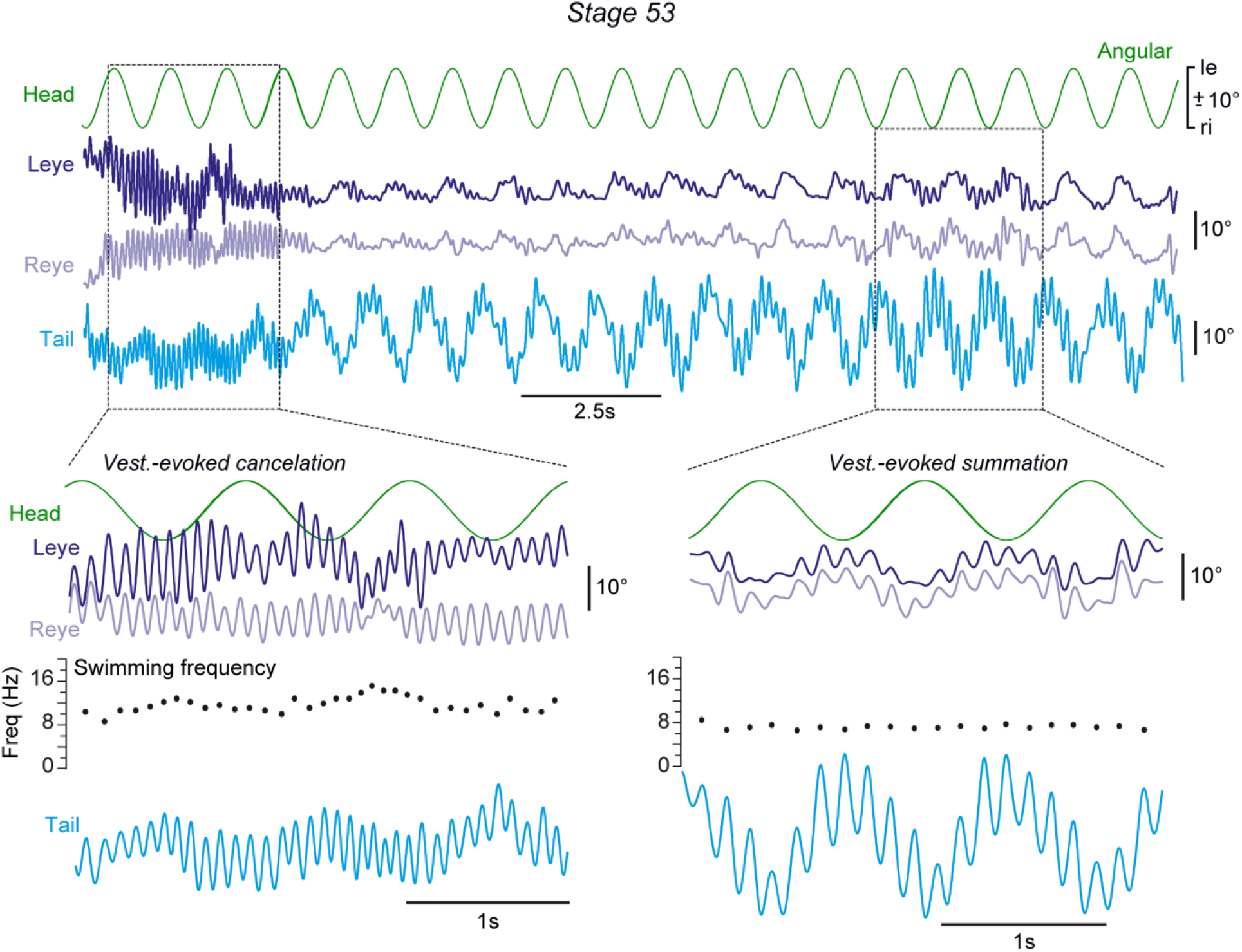
Computational interactions between vestibulo-ocular and spino-ocular motor signals depends on swimming frequency. Movements of the left (Leye, dark blue) and right eye (Reye, light blue) during horizontal sinusoidal head rotation and swim-related tail undulations; in the same preparation, horizontal semicircular canal signals are cancelled (inset at lower left) at a tail undulatory swim frequency (Freq.) of ∼10-12 Hz but are summated with locomotor efference copy signals (inset at lower right) when swim-related tail undulations occur at ∼6 Hz.

